# Zika Virus: Endemic Versus Epidemic Dynamics and Implications for Disease Spread in the Americas

**DOI:** 10.1101/041897

**Authors:** Sharon Bewick, William F. Fagan, Justin Calabrese, Folashade Agusto

**Affiliations:** Department of Biology, University of Maryland, College Park, MD 20742; Conservation Ecology Center, Smithsonian Conservation Biology Institute, Front Royal, VA 22630; Department of Ecology and Evolutionary Biology, University of Kansas State, Lawrence, KS 66045

## Abstract

Since being introduced into Brazil in 2014, Zika virus (ZIKV) has spread explosively across Central and South America. Although the symptoms of ZIKV are generally mild, recent evidence suggests a relationship between prenatal exposure to ZIKV and microcephaly. This has led to widespread panic, including travel alerts and warnings to avoid pregnancy. Because ZIKV is an emerging disease, response efforts are complicated by limited understanding of disease dynamics. To this end, we develop a novel state- and class-structured compartment model for ZIKV. Our model shows that the risk of prenatal ZIKV exposure should decrease dramatically following the initial wave of disease, reaching almost undetectable levels in endemic systems. Our model also suggests that, depending on ZIVK transmission levels in the Americas, efforts to reduce ZIKV prenatal exposures through mosquito management and avoidance may have minimal benefit, and may even result in increased risk of microcephaly in later years of an outbreak.

---

After being discovered in Ugandan forests in 1947^1^, Zika virus (ZIKV) remained a relatively minor arboviral disease for 60 years^2^. In 2007, however, an outbreak of ZIKV on Yap Island^3^ in the Pacific Ocean signaled spread of the virus beyond its historic range^2-4^. From Yap Island, ZIKV was transported to French Polynesia in 2013^5^ and then on to Brazil in 2014^6-8^. Once in Brazil, the virus took off, ‘spreading explosively’^9^ throughout both South and Central America. By early 2016, for example, local transmission of ZIKV had been reported in 20 countries and territories in the Americas^10^. Initially, ZIKV was not viewed as a significant public health threat. Indeed, with a negligible mortality rate and symptoms resembling a mild form of dengue (DENV)^2^, the ZIKV outbreak appeared to be more of a nuisance than a public health emergency. In November 2015, however, alarms were raised about a potential connection between ZIKV transmission and increasing rates of newborn microcephaly^11^.

Currently, the link between ZIKV and microcephaly is only postulated, not proven^12^. Nevertheless, the >20-fold increase in microcephaly in regions of Brazil where ZIKV is spreading^13^ has been enough to initiate drastic precautionary action. The United States Centers for Disease Control (CDC), for example, posted a travel alert recommending that pregnant women avoid regions in the Caribbean and Latin America where ZIKV transmission is ongoing^14^. Meanwhile, public health officials in El Salvador and Colombia have suggested that women delay pregnancy up to two years until ZIKV outbreaks can be controlled^15^.

Like other viruses in the genus *Flavivirus*, for example DENV, West Nile Virus (WNV) and Yellow Fever Virus (YFV), ZIKV is primarily spread by mosquitoes. For ZIKV, the main vectors appear to be members of the genus *Aedes*^16^, including the notorious *Ae. aegypti*. This is of concern because *Aedes* species are widespread in warmer temperate and tropical regions^17,18^. In addition, although chemical larvicides and adulticides are somewhat effective at reducing certain *Aedes* populations, these mosquitoes can reproduce in very small containers of standing water. This makes complete eradication difficult^19^, casting doubt on claims that mosquitoes can be controlled sufficiently to stop ZIKV transmission.

One puzzling aspect of the recent ZIKV outbreak in South/Central America is why correlation is only now emerging between prenatal exposure to ZIKV and microcephaly. ZIKV is not a particularly new disease. In fact, phylogenetic analyses indicate that ZIKV has likely been circulating in Africa and parts of Asia for approximately 100 years^20^. Why, then, has microcephaly not been reported in African/Asian countries at the same alarming rates currently making headlines in Brazil? Assuming that there is a link between ZIKV and microcephaly, there are several potential explanations. First, there may be under-reporting of microcephaly in Africa/Asia, making tracking of microcephaly difficult and complicating comparison to the situation in South/Central America. Second, the ZIKV strain currently circulating in the Americas, which is derived from a more recently evolved Asian lineage^8,21^, might be associated with more severe complications, including microcephaly. Third, intrinsic differences may exist between African/Asian and American populations, and this might modify either the extent of ZIKV spread or the risk of severe ZIKV complications. Population-wide differences could, for example, reflect genetic predisposition^22^. Alternatively, differences in microcephaly incidence may indicate differing immunological statuses of people in the two regions, and this might be a function of previous exposure to ZIKV or other related diseases.

Without knowing why disease epidemiology differs between South/Central America versus Africa/Asia, it is difficult to predict how the ZIKV outbreak will progress. Epidemiological modeling is a powerful tool that has previously proven useful for understanding the spread and dynamics of other vector-borne diseases, including other flaviviruses^23,24^. Here, we construct a compartment model (see Supplemental Information I, Figure S1) to describe ZIKV transmission, both in countries where the disease is endemic, and in countries where the disease has been newly introduced. While pregnancy related complications of viral diseases have been studied previously, particularly in the case of rubella ^25,26^, this is a new feature within vector-borne diseases. Consequently, understanding the risks of prenatal ZIKV exposures requires extending traditional vector-borne disease models to a novel age-and class-structured framework that allows us to focus directly on ZIKV infection during pregnancy.

## Results

Because ZIKV is an emerging disease, very little is known about its biology. Consequently, we define our model based on the related flavirus DENV. However, to provide additional information, we also consider ZIKV seroprevalence data from regions where ZIKV has been present for many years. This allows us to identify more likely ZIKV parameter ranges within the broader ranges derived from DENV studies. Specifically, we consider a ‘low transmission’ scenario, an ‘intermediate transmission’ scenario and a ‘high transmission’ scenario. The first two scenarios are determined, respectively, by fitting model predictions to juvenile seroprevalence data and overall (age-adjusted) seroprevalence data from a study in Kenya^27^. The ‘high transmission’ scenario is based on average values from the ranges predicted for DENV (see Supplemental Information II). Notably, our intermediate transmission scenario not only matches endemic seroprevalence rates in Kenya, but also matches epidemic (one year into an outbreak) seroprevalence rates in Micronesia^28^ (see Supplemental Information II).

### High Rates of ZIKV Infections in Pregnant Women Early in an Epidemic

Figure 1 shows the predicted number of women who will experience a ZIKV infection during pregnancy as a function of the number of years since ZIKV arrival in a region (intermediate transmission scenario, see Supplementary Information II). Similar dynamics are observed for all but the most conservative estimates of ZIKV transmission, suggesting that one third to one half of women who are or become pregnant during the first year of a ZIKV outbreak will experience a ZIKV infection at some point during their pregnancy (see Supplementary Information II Figure S3 and Supplementary Information III Figure S4). Assuming that there is a link between ZIKV and microcephaly, this explains the dramatic increase in microcephaly rates in Brazil. Indeed, even if ZIKV crosses the placenta in only a small fraction of infections, or only affects the fetus during the early stages of pregnancy (see also Supplementary Information IV, Figure S5) this will still result in a sizeable fraction of babies born with ZIKV complications.

**Figure 1.**
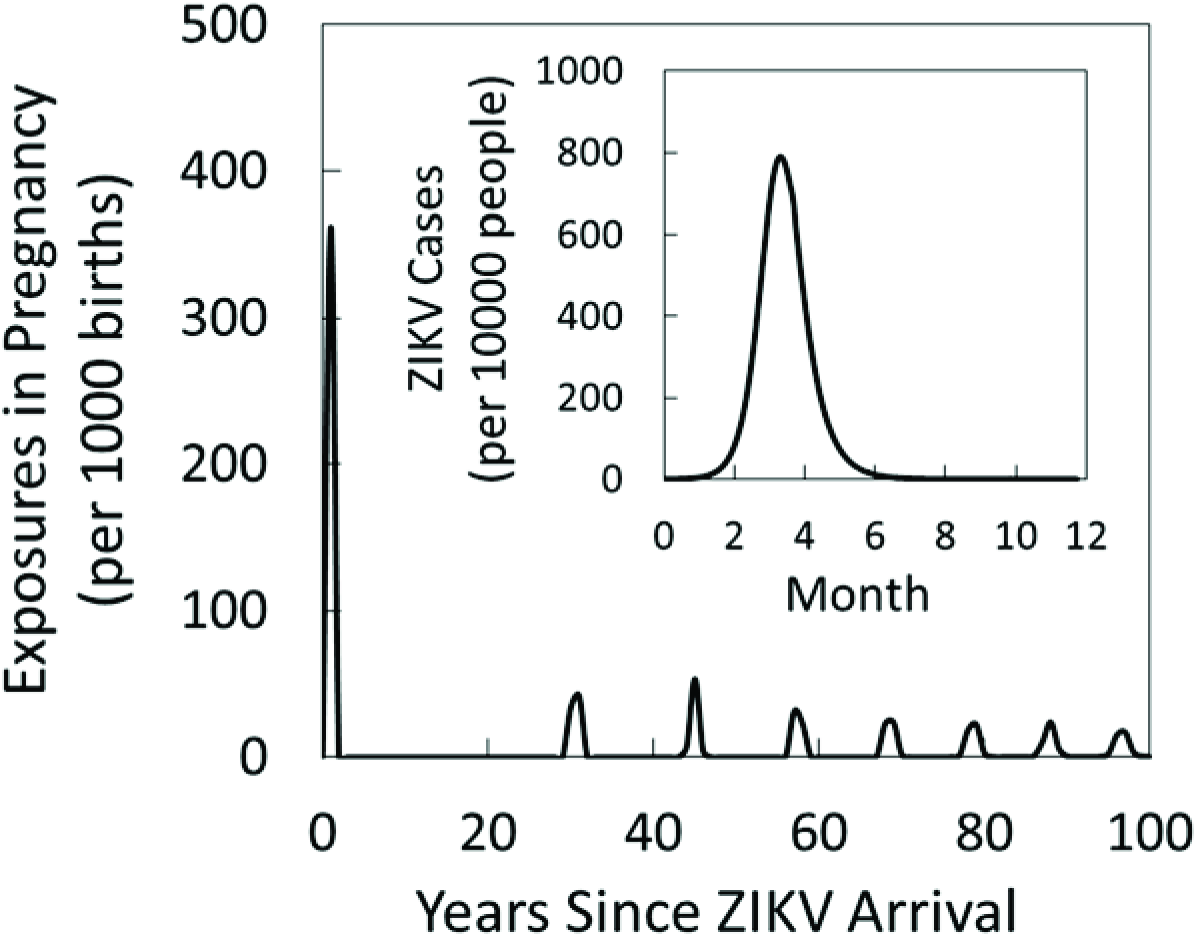
Predicted ZIKV dynamics, showing the number of women who experience a ZIKV infection during pregnancy as a function of the number of years since ZIKV arrival in the country or region. For this figure, we assume our intermediate transmission scenario (see Supplemental Information II). The inset shows the total number of ZIKV cases during the first year of the epidemic.

Alarmingly, efforts to minimize ZIKV exposure rates may have limited benefits. In Figure 2, for example, we see that, at least under the high transmission scenario (black lines, see Supplemental Information II), ZIKV infections in women pregnant during the first year of a ZIKV epidemic are nearly independent of mosquito biting rates, mosquito recruitment rates and mosquito death rates, which are the key targets of insect repellant, larvicides and adulticides respectively. For the intermediate transmission scenario (dark grey line), control efforts are more effective, although reducing exposures in pregnant women to below 1% still requires a 30% reduction in mosquito biting rates, a 52% reduction in mosquito recruitment or a 45% increase in mosquito death rates (or some combination thereof). For the low transmission scenario (light grey line), ZIKV is much closer to the threshold necessary for disease persistence (*R*_0_ = 1). As a consequence the epidemic is much easier to bring under control, making mosquito management efforts more effective.

**Figure 2.**
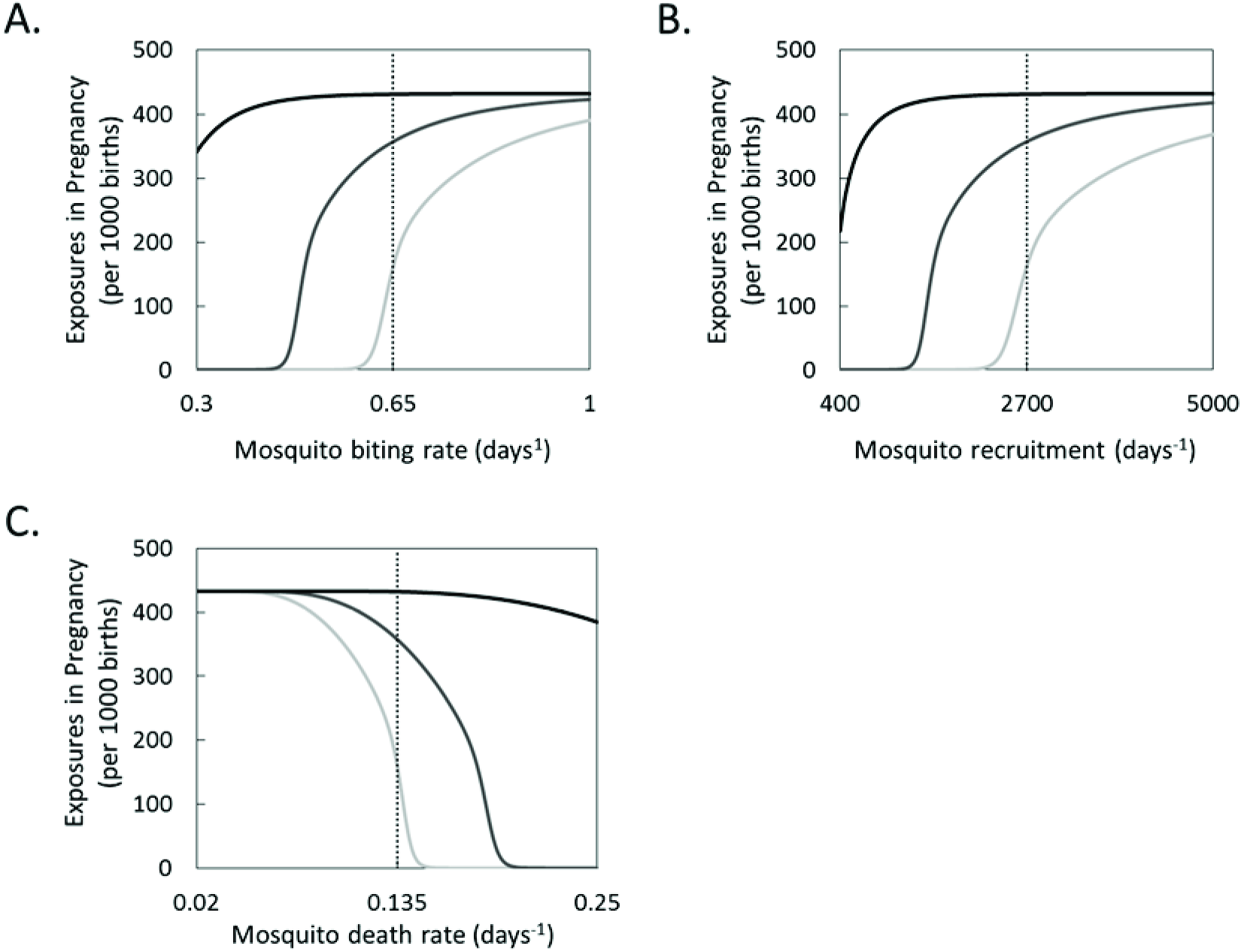
The number of pregnant women exposed to ZIKV during the first year of an outbreak as a function of (A) mosquito biting rate, (B) mosquito recruitment and (C) mosquito death rate assuming low (*β*_*v*_ · *β*_*h*_ = 0.0442 light grey), intermediate (*β*_*v*_ · *β*_*h*_ = 0.0763 dark grey) and high (*β*_*v*_ · *β*_*h*_ = 0.319 black) ZIKV transmission scenarios (see Supplemental Information II). Vertical dotted lines represent average values for the parameters under consideration, and thus can be thought of as starting points for control efforts.

### Dramatic Decrease in Prenatal ZIKV Exposures Within 1-2 Years

Despite the disheartening statistics for the first year of a ZIKV epidemic, there is a silver lining. In the years following the initial dramatic explosion of ZIKV cases, we predict a sudden and rapid decrease in prenatal ZIKV exposures, ultimately reaching an almost undetectable level. Importantly this decrease is not a result of any control strategies or vector management efforts. Instead, it is an intrinsic property of the system - a property that arises as a result of the interplay between transmission of infection, build-up of immunity in the human population, and the timing of human reproduction. In fact, in an ironic twist, efforts to control ZIKV transmission through mosquito management and avoidance of biting insects may actually backfire (at least in the long run), increasing, rather than decreasing the ultimate number of ZIKV prenatal exposures. This can be seen explicitly in Figure 3. Here we show the number of years it takes for the cumulative ZIKV caseload in pregnant women in systems with mosquito control (i.e. reduced mosquito biting rates through repellant use, reduced mosquito recruitment rates through use of larvicides and increased mosquito death rates through use of adulticides) to surpasses the same caseload in systems without mosquito control. As expected, cross over points occur earlier in systems with higher transmission, and do not occur at all in systems with very low transmission. Consequently, whether or not mosquito control is the best long term strategy for preventing ZIKV infections in pregnant women depends on both the natural rate of ZIKV transmission, and also the efficacy of control strategies. If ZIKV transmission is naturally relatively low and control efforts are relatively effective, then the use of repellants, larvicides and adulticides can result in both immediate and long-term reductions in ZIKV infections in pregnant women. However, if transmission is naturally higher, and control efforts less effective, then the use of repellants, larvicides and adulticides, while preventing ZIKV infections in pregnant women over the near term, can actually result in higher exposure rates in pregnant women over longer time windows.

**Figure 3.**
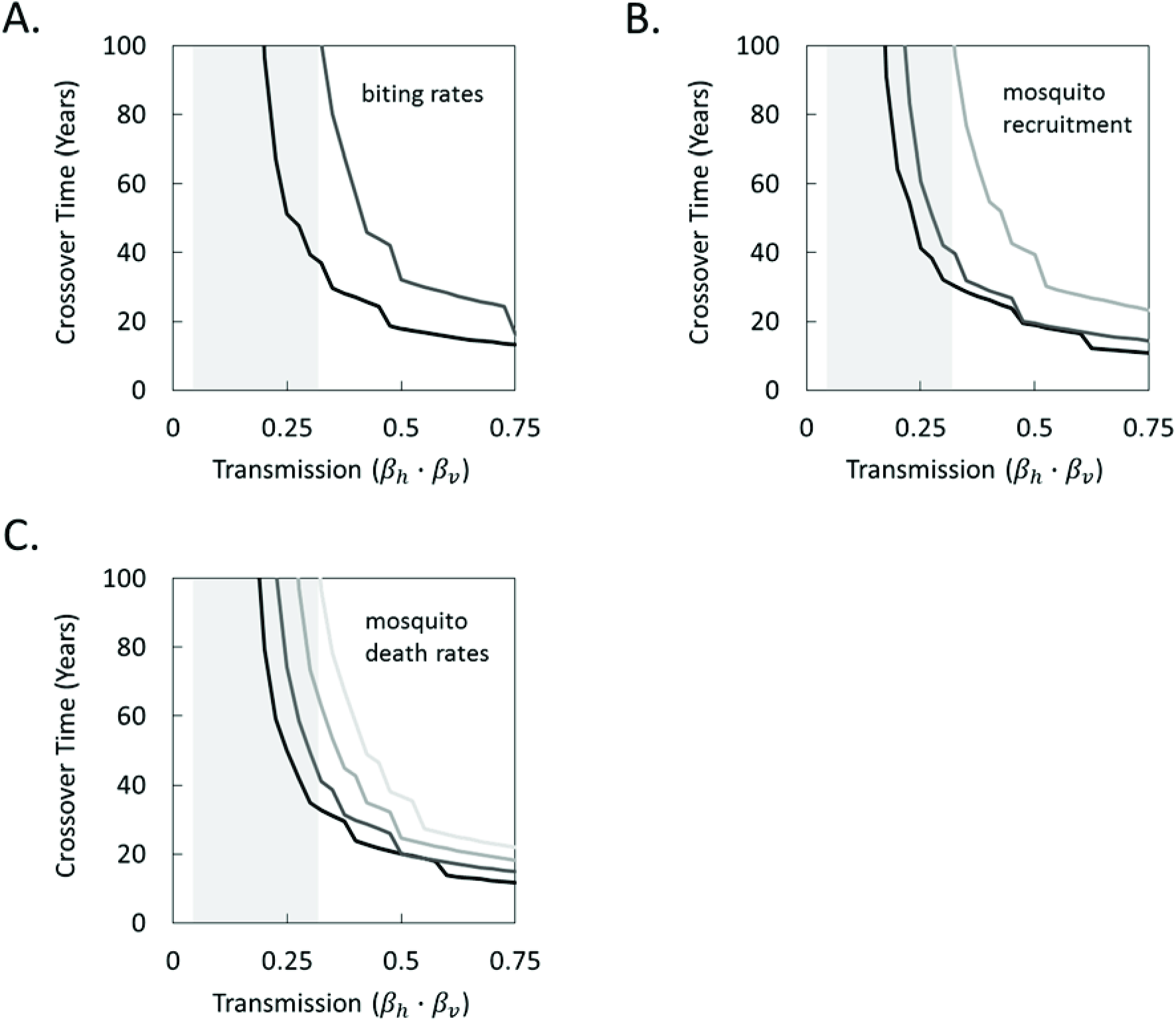
Plots showing the number of years before the cumulative ZIKV caseload among pregnant women is higher (crossover time) as a result of control efforts using (A) reductions in biting rates, (B) reductions in mosquito recruitment and (C) an increase in mosquito death rates. Reductions (increases) are as follows: 25% (black), 50% (dark grey), 75% (medium grey) and 100% (light grey). Reductions not shown in panels (A,B) are sufficient to ensure that the cumulative ZIKV caseload in pregnant women is lower in the system with control for at least 100 years. Crossover times are considered across the entire range of potential transmission probabilities based on DENV models (see Supplemental Information II, Table S1); grey boxes are used to denote narrower ranges based on our three transmission scenarios (low, intermediate and high).

### High Levels of Disease Transmission Prevent Prenatal Exposures

Figure 4 expands on Figures 2 and 3, showing yearly prenatal exposures to ZIKV that would be expected in regions where the virus has been endemic for many years (i.e., equilibrium exposure rates). Compared to the 30-40% of pregnant women infected during the first year of a ZIKV epidemic (see Figure 1, Supplemental Information II, Figure S3), predicted yearly exposures in regions where the disease is endemic are much lower (see also Supplemental Information III, Figure S4), typically below 5 infections per 1000 births. Similar to epidemic predictions, analysis of endemic systems suggests that, particularly for the high transmission scenario, prenatal ZIKV exposures decrease with mosquito biting and recruitment rates and with mosquito life expectancies. Again, this is a result of the interplay between disease transmission, build-up of population-level immunity, and the timing of reproduction. In particular, when mosquito biting rates, recruitment, and longevity are high, so too is disease spread. As a result, there is ample opportunity to acquire a ZIKV infection, meaning that very few individuals born in a region with endemic ZIKV will reach reproductive age without having been previously exposed to the virus (see dashed lines in Figure 4). However, for lower mosquito biting rates, recruitment, and life expectancies, opportunities for disease acquisition are reduced. If this reduction is not sufficient to make the likelihood of infection during pregnancy negligible, then the net result can be a higher risk of disease acquisition while pregnant, despite a lower overall risk of disease acquisition at any stage of life.

**Figure 4.**
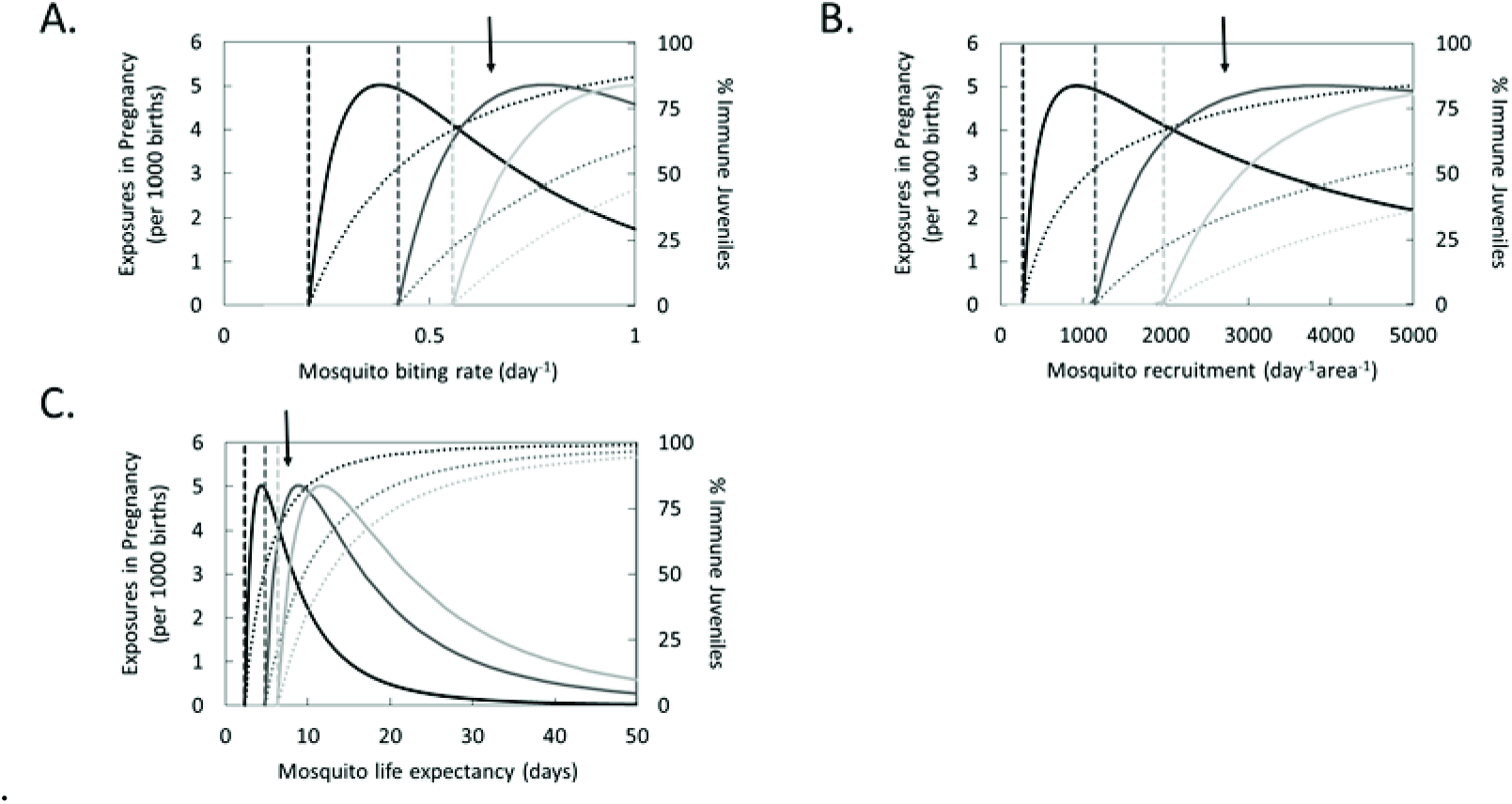
Number of women who experience an active ZIKV infection at any point during pregnancy (solid lines) and percentage of children (below reproductive age) that have acquired ZIKV immunity (dotted lines) as a function of (A) mosquito biting rates, (B) mosquito recruitment rates and (C) mosquito life expectancy in a region with endemic disease (i.e., a system at equilibrium). Results are shown for high (black), intermediate (dark grey) and low (light grey) ZIKV transmission scenarios. The dashed vertical lines indicate the mosquito biting rates, recruitment rates and life expectancies that give *R*_0_ = 1 (i.e., the threshold for disease persistence). The black arrows show the average values of the parameters under consideration (see Supplemental Information II, Table S1), and thus can be thought of as starting points for control.

### Model Parameter Exploration

To explore the implications of our model findings over the full range of parameter space (see Table S1), we use a Latin Hypercube Sampling (LHS) analysis (see Materials and Methods), yielding Figure 5. Comparing partial rank correlation coefficients (PRCCs) for endemic versus epidemic ZIKV, we find that the majority of parameters that are negatively correlated with prenatal exposures in endemic regions are positively correlated with prenatal exposures in regions where ZIKV has been newly introduced (i.e., is still epidemic). Thus, consistent with Figures 1-3, mosquito biting rates, recruitment, and life expectancy are associated with a reduced risk of prenatal exposures when ZIKV circulation reaches an equilibrium level, but can result in an increased risk of prenatal exposures during the initial wave of disease. Not surprisingly, parameters associated with a high risk of prenatal exposures under epidemic conditions are the same parameters associated with a large basic reproduction number, *R*_0_ (i.e., they tend to make disease spread more favorable, see Materials and Methods). These are also the same parameters associated with high rates of ZIKV immunity in children. This latter finding reiterates the mechanism responsible for the counterintuitive reduction in prenatal exposures when there is high disease transmission in endemic systems. In particular, when ZIKV has been in a region for a number of years, parameters positively correlated with childhood immunity ensure that the majority of children acquire ZIKV prior to reaching reproductive age.

**Figure 5.**
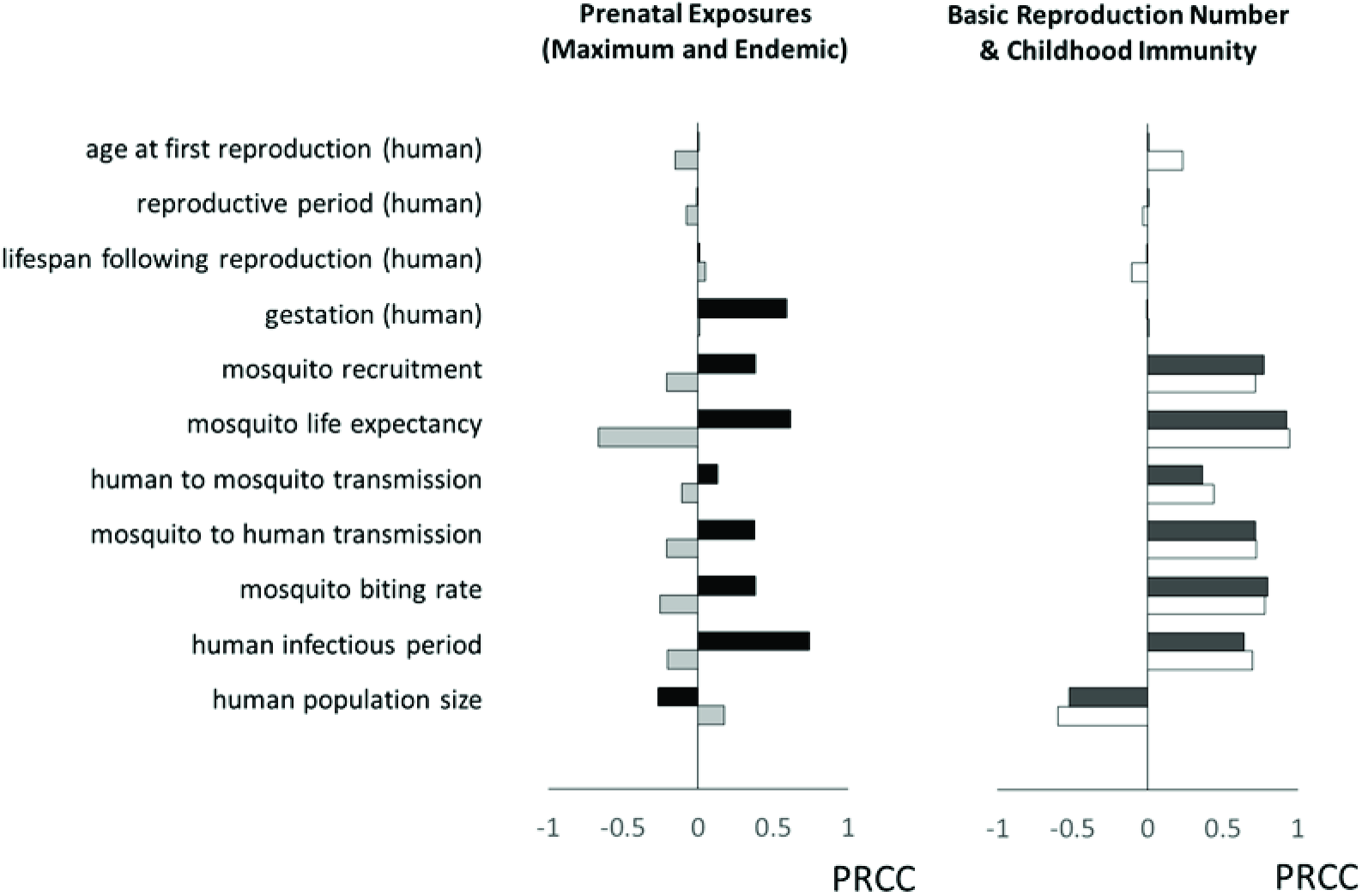
Partial Rank Correlation Coefficients (PRCCs) for the fraction of women who experience a ZIKV infection during pregnancy at the peak of an outbreak (black), and after ZIKV transmission has reached endemic levels (light grey), as well as PRCCs for the ZIKV basic reproduction number (white) and the fraction of children who acquire ZIKV immunity prior to reproductive age in regions where ZIKV is endemic (dark grey). For infections during pregnancy and childhood immunity, we use the parameter ranges in Table S1, but restrict the following: *b* ∈ (0.4, 1), *A* ∈ (1000,5000) and *β*_*h*_ ∈ (0.15, 0.75) to avoid issues with non-monotonicity. For the basic reproduction number, we use the full ranges for all parameters.

## Discussion

Since the announcement that microcephaly may be related to ZIKV infection during pregnancy, public health officials have been scrambling to find solutions for halting the current ZIKV outbreak in South and Central America. Meanwhile, North America is preparing for the inevitable arrival of ZIKV, which is expected in the coming months, as temperatures begin to warm and mosquitoes become active. Unfortunately, without a full understanding of disease dynamics, it is difficult to predict how the future course of the ZIKV epidemic will unfold, and nearly impossible to determine the best strategies for managing it. In this paper, we take a step towards understanding the population level consequences of ZIKV transmission by constructing a dynamic compartment model for ZIKV. ZIKV is unique among vector borne viruses in that it poses its greatest threat during pregnancy. Consequently, we build an age-and class-structured model that allows us to specifically consider the risk of prenatal exposure to the virus.

### Microcephaly in Brazil

Interestingly, our model predictions immediately reconcile the strikingly high incidence of microcephaly in Brazil, despite this never being documented as a complication of ZIKV in Africa or Asia, where the disease has been endemic for many years. In particular, we find that rates of infection of pregnant women during the first one to two years of a ZIKV outbreak should be remarkably high (typically 30-40%, see Figure 1 and Supplemental Information III Figure S4). However, after this initial surge, there should be a precipitous decline in ZIKV infections during pregnancy and, by the time the disease has become endemic, very few pregnant women should acquire ZIKV. The cause is two-fold. First, after the initial wave of infection, the overall disease burden in the community decreases as immunity builds up in a previously naïve population. Second, prenatal exposures are further suppressed by the age structure associated with human reproduction. In particular, because most females do not reproduce until they are at least 15 years old, and usually older^29^, there is sufficient time during childhood to acquire ZIKV and develop immunity. This immunity then protects the fetus, even if the mother is bitten by an infectious vector. In other words, high rates of ZIKV infection during childhood act as a ‘natural vaccine’ that effectively prevents ZIKV infection during the critical pregnancy stage.

Notice that the protective effects of natural ZIKV infection hinge upon having high levels of ZIKV circulating in the community. This, in turn, depends upon mosquito densities and biting rates, as well as the intrinsic ability of ZIKV to spread from humans to mosquitoes and vice versa. Although dengue is thought to exhibit the type of ‘flash and fade’ dynamics required for the protective mechanisms we propose^30^, the precise rates of transmission of ZIKV are unknown. Nevertheless, there are several lines of evidence to suggest that ZIKV also exhibits the transmissibility necessary to achieve high population-level immunity. First, the explosive dynamics of ZIKV in both French Polynesia^5^ and Brazil^11^ indicate that, once introduced into a population, ZIKV spreads rapidly. Second, there have been a number of estimates of ZIKV prevalence based on serological studies and surveys, all of which suggest relatively high rates of coverage. The first study, in Kenya in 1970, indicated that an average of 52% (not age-adjusted) of the human population in the Malindi District had ZIKV antibodies, though only 1% and 3% of the populations were covered in Central Nyanza and the Kitui District respectively, where *Aedes aegypti* are largely absent^27^. The second study, in Nigeria in 1979, again suggested that 52% (not age-adjusted) of the population was protected against ZIKV^31^. The third study, also in Nigeria in 1980, indicated that 56% (not age-adjusted) of the population had ZIKV antibodies^32^. More recently, a study on Yap Island following the 2007 outbreak, indicated that approximately 73% of the population had been exposed to ZIKV during the epidemic^4^. Likewise, in outbreaks in Micronesia and Polynesia, an estimated 80% and 50% of the population was exposed to ZIKV respectively^28^. These percentages are similar to estimates from our model (see Figure 4, Supplemental Information II), and should be sufficient to observe the protective benefits against infection during pregnancy. Similarly supporting our model predictions, in both Micronesia and Polynesia, outbreaks lasted approximately 5 months, which is very close to the length of the outbreak predicted by our model (see Figure 1, Supplemental Information II Figure S3).

### Management Implications

The issue of ‘natural vaccination’ raises an interesting but difficult question. Is the best approach to managing ZIKV infection really to target mosquitoes and mosquito exposure? Although such targeting might help to prevent near-term infection of pregnant woman (see Figures 2), it also slows the rate at which ZIKV immunity is acquired in the population as a whole. This delay means that women who would have been exposed to ZIKV prior to pregnancy under high transmission scenarios, might not come in contact with the disease until they are carrying a child under lower transmission scenarios. This is exactly what we see in model predictions – over multiple decades, efforts to control mosquito biting rates, mosquito recruitment rates and mosquito death rates can actually result in higher endemic rates and more cumulative cases of ZIKV in pregnant women (see Figures 3-4), particularly if ZIKV transmission is naturally very high, making control efforts less effective (see Figure 2).

Notice that the argument against mosquito control is similar to arguments that were made decades ago about the benefits and consequences of rubella vaccination^26^. Ultimately, whether management is the best strategy depends heavily on natural ZIKV transmission rates. Consequently, we want to stress that we in no way support immediate cessation of mosquito control programs in South and Central America. Before such a decision can be made, more information is required. This includes laboratory measurements of rates of ZIKV transmission to and from mosquitoes (see Supplemental Information II), measurements of mosquito densities in affected regions, extensive surveys of ZIKV seroprevalence rates (including in the Americas and among pregnant women), determination of the pregnancy stages associated with risk, and longer term analysis of waning immunity (which could compromise some of the protective benefits that we predict from disease acquisition, see below). Another, separate consideration is the effectiveness of mosquito control programs. If mosquito control can successfully reduce ZIKV transmission levels to near or below what is necessary for disease persistence (*R*_0_ <1, see Materials and Methods), then mosquito eradication is unquestionably the best strategy (see Figure 4).

Interestingly, as compared to the uncertain consequences of mosquito control, the benefits of delayed pregnancy are much clearer. Indeed, we predict that, in regions with intense mosquito activity, high rates of prenatal ZIKV exposure should only last for one or two years. Therefore, even if it proves impossible to reduce mosquito populations sufficiently to stop or dramatically slow the spread of ZIKV, there may still be a benefit to postponing pregnancy. Consequently, while the recommendations to delay pregnancy by the El Salvadorian and Colombian governments were likely motivated by an overly optimistic expectation that ZIKV transmission can be halted, this advice may nonetheless be highly effective for reducing rates of microcephaly. Delayed pregnancy has the advantage of avoiding ZIKV complications in the near-term, while still allowing rampant ZIKV spread that then ‘vaccinates’ females, protecting them for when they do eventually become pregnant. An even better, though more costly, solution would be to administer serological tests to women who want to become pregnant. Women with ZIKV antibodies could proceed with pregnancy plans 1-2 months following a positive test, while women who have not been infected could be advised to wait for an additional period of time and then retest.

### Model Limitations

The main findings of our model – that the risk of ZIKV infection in pregnant women should decrease dramatically in 1-2 years, and that long-term ZIKV infections in pregnant women should be extremely low, are highly robust to model parameters (see Supplemental Information III Figure S4). However, there are several structural assumptions that, though very likely based on findings from other flaviviruses, may compromise our results. First, and most importantly, we assume life-long immunity. Waning immunity and, in particular, immunity that lasts < 30 years would allow women infected during childhood to become susceptible again prior to or during their reproductive period. Likewise, if a large fraction of people do not develop immunity at all, this could also negate our predictions. Evidence for high numbers of people experiencing ZIKV re-infections would thus bring into question the ZIKV dynamics that we predict. Another assumption of our model is that, once recovered, people remain recovered. Given, however, that ZIKV can infect immune privileged sites like the brain and reproductive tissues, ZIKV resurgence months after the initial exposure is a possibility. Depending on rates of resurgence and also the periods between initial infection and resurgence, this could potentially alter both transmission dynamics and the risk of experiencing an active infection during pregnancy. Again, evidence of people experiencing multiple, active ZIKV infections could bring into question our predicted dynamics. On a related note, evidence of women giving birth to more than one child (excluding twins, etc.) with microcephaly could indicate a violation of some of our assumptions, since this might indicate re-infection, resurgence or even that there is a risk to the fetus even after the mother has cleared the infection. Finally, our model assumes a non-spatial, well-mixed population. As a result, it should only be interpreted as explaining local ZIKV dynamics. This means that, although we do not expect extended periods of ZIKV spread in any one location, it could easily take many years for ZIKV to spread across the Americas. Furthermore, ZIKV infection will likely continue to be a serious problem for travelers to the Americas for many years.

### Conclusions

In the coming months, we anticipate intense study of ZIKV. Hopefully, this will provide researchers with increased information on the virus’ natural history and epidemiology. As this information becomes available, it will enable progressively more detailed models of ZIKV transmission that will allow for clearer predictions of how the ZIKV outbreak is likely to impact the Americas. More accurate estimates of disease transmission rates, incorporation of latency periods in vector and host populations, the potential for waning immunity and even seasonality effects are some of the details that continued study should elucidate. This paper presents a framework for including all of these anticipated effects. New information will help to tighten model predictions, including better estimation of whether ZIKV can be sufficiently suppressed by mosquito control techniques and, if not, how the trade-offs between short-term mosquito control and long-term population immunity should be balanced to minimize the risks of prenatal ZIKV exposure. Ultimately, this will be valuable information for management of what is surely one of the most pressing disease outbreaks in recent history.

## METHODS

We build a dynamic compartment model for ZIKV (see Supplemental Information, Figure S1) and use this to analyze disease dynamics, both in regions where ZIKV did not previously exist, and in regions where the virus is endemic. Because the primary concern with this virus is its potential to cause microcephaly in newborns, we focus on a structured population model that accounts for age-structure, gender, and pregnancy status in the human population. For human disease transmission, we use an S-I-R (Susceptible-Infected-Recovered) framework that assumes life-long immunity in people who have had a ZIKV infection. We use an S-I-R model based on observations of neutralizing ZIKV antibodies in patients that have recovered from ZIKV^31^. Lifelong immunity is also a common attribute for other closely related flaviviruses, for example DENV^30^ and YFV^33^. Specifically, our model is as follows:

### Humans (S-I-R)

*Pre-reproductive females and males*:

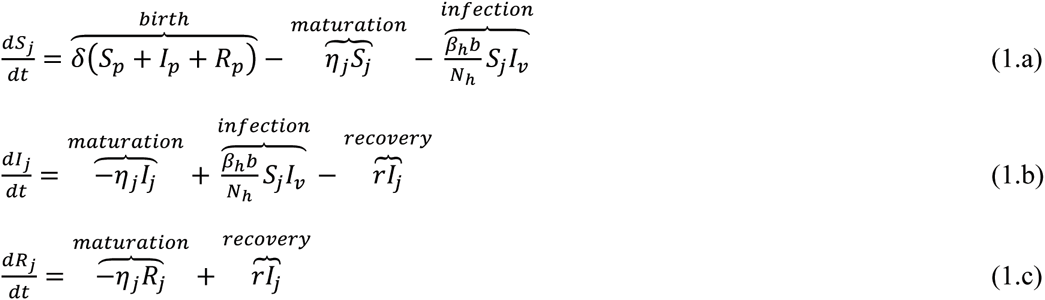

*Reproductive females (not pregnant)*:

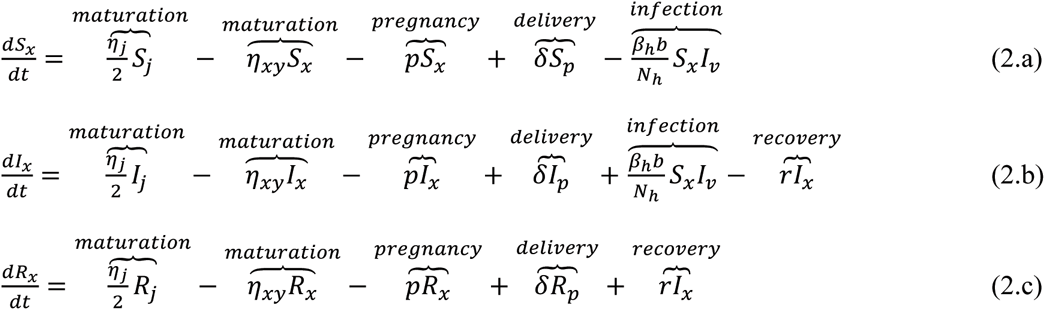

*Reproductive females (pregnant)*:

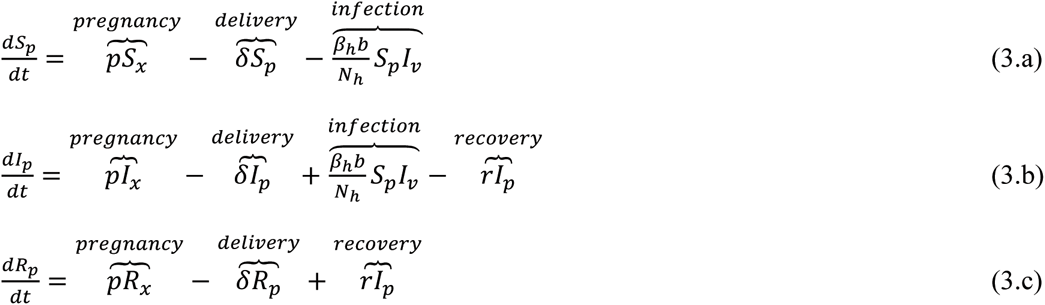

*Reproductive males*:

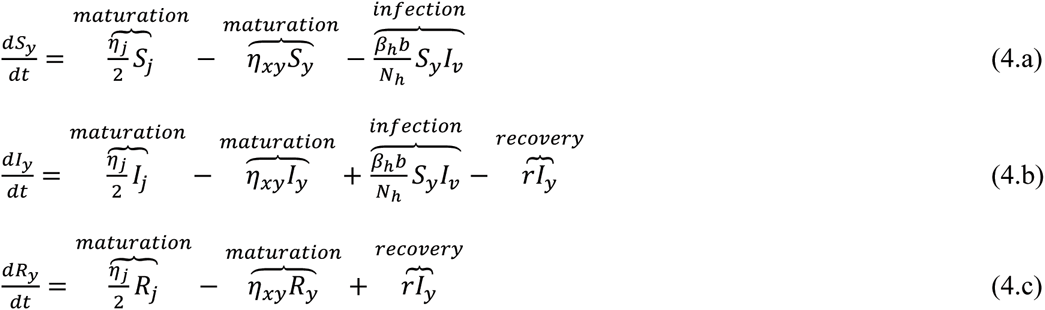

*Post-reproductive females and males*:

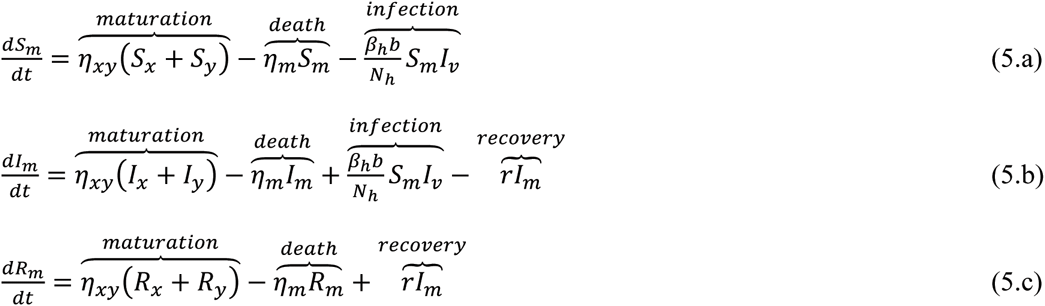

where *S, I* and *R* are populations that are susceptible to, infected with, and recovered from (and thus immune to) ZIKV respectively and subscripts on the state variables are: *j* for children, *x* for reproductive-aged females that are not pregnant, *p* for reproductive-aged females that are pregnant, *y* for males within the age range of reproductive females, and *m* for adults beyond reproductive age (as based on female reproduction). Notice that, in equations (1-5), mortality only occurs in the post-reproductive class. Although this is not, strictly speaking, true, it is a good approximation for countries where the primary source or mortality is senescence. We also do not assume additional death in infected classes because, unlike many other flaviviruses, mortality associated with ZIKV is negligible (at least, outside of the prenatal stage).

For the vector population, we assume an S-I model. Specifically:

***Mosquitoes (S-I):***

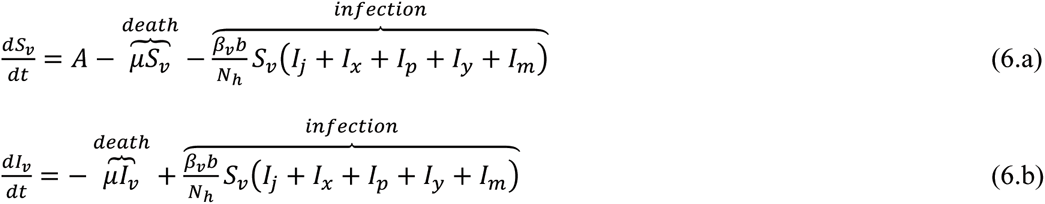

where *S*_*v*_ are the susceptible vectors and *I*_*v*_ are the infected vectors. We choose this model for the vector because it is structurally identical to a number of classic DENV models^23^. Furthermore, it is simple enough to allow us to focus on the effects of age-structure and pregnancy in the human population and to avoid issues of model over-parameterization that would diminish the model’s predictive value. In particular, notice that, although we have 17 state variables, there are only 12 parameters (11 if you assume a constant human population size), most of which are well defined based on human life-history. For mosquito parameters, we use values previously determined from analysis of DENV models. We believe that this is valid, since DENV is also spread by *Aedes* mosquitoes, and *Ae. aegypti* in particular. For disease parameters (the probability of mosquito to human transmission, *β*_*h*_, the probability of human to mosquito transmission, *β*_*v*_, and the human infectious period, 1/*r*), we again select ranges based on DENV. For the human infectious period, the parameter range for DENV is consistent with preliminary estimates from recent ZIKV outbreaks^34^. For transmission probabilities, DENV parameter ranges are broad. We thus use seroprevalence data from ZIKV studies to further inform likely ranges for these parameters (see Supplemental Information II).

*Basic Reproduction Number*: We find the basic reproduction number, *R*_0_, for the system in equations (1-6) as follows^35^:

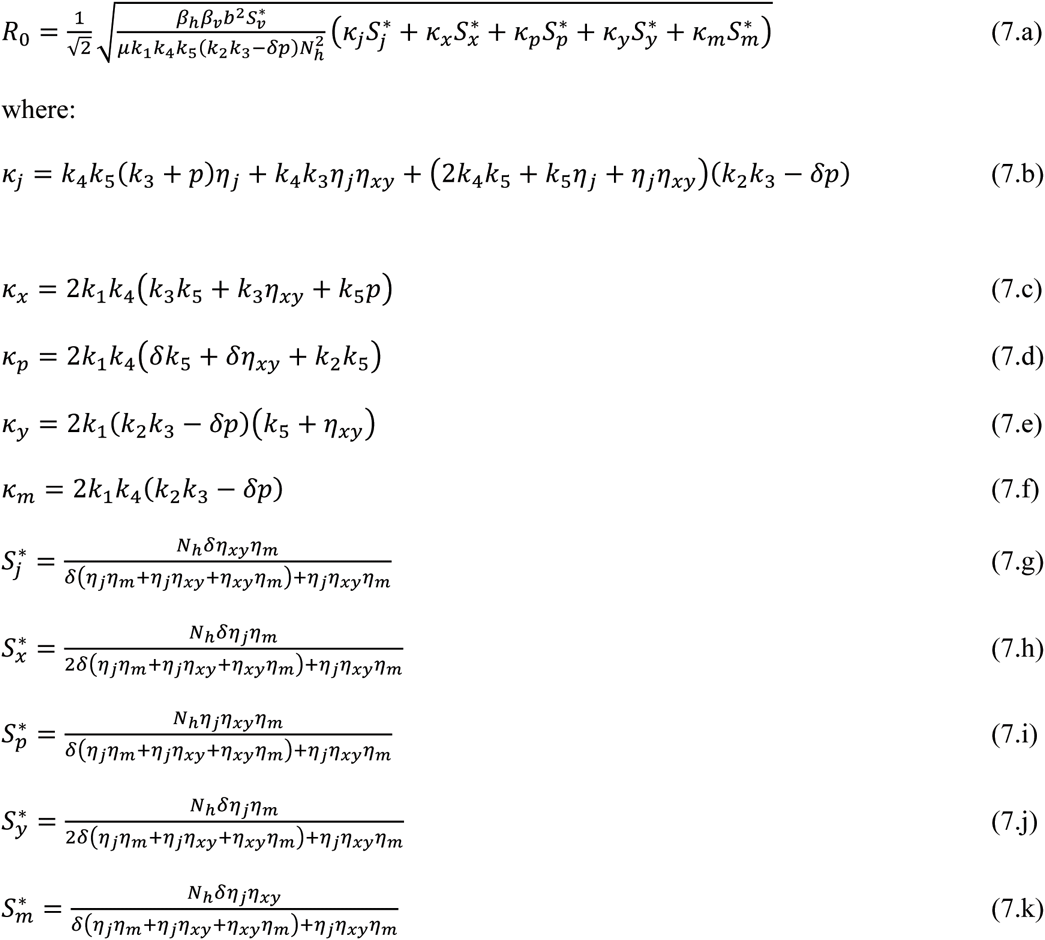

and *k*_1_ = *η*_*j*_+*r, k*_2_ = *η*_*xy*_ + *p* + *r,k*_3_ = *δ* + *r,k*_4_ = *η*_*xy*_ + *r*, and *k*_5_ = *η*_*m*_ + *r*. Notice that equations (7.g-k) assume a fixed population size, thus *p* = 2*η*_*xy*_. When *R*_0_ > 1 the disease-free equilibrium is unstable, meaning that ZIKV spread is predicted. In contrast, when *R*_0_ < 1, the disease-free equilibrium is stable, and ZIKV should not persist in the system.

*Infection During Pregnancy:* Focusing on the risk of microcephaly, the fraction of women who experienced an active ZIKV infection at any point during a pregnancy in year *n* is:

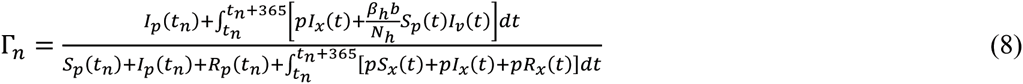

where *t* is measured in days and *t*_*n*_ is the first day of year *n*. Notice that this means women whose pregnancies (infections) span two years (e.g., begin in October and end in June) are counted towards pregnancy (infection) totals for both years. Throughout the paper, we use equation (8) as a marker of disease intensity and the cost of ZIKV transmission. Specifically we explore how the risk of ZIKV infection during pregnancy changes depending on the length of time that ZIKV has been present in a country, as well as vector biology and/or management actions. We then compare the risk of acquiring ZIKV during pregnancy in a country where the virus is endemic with the same risk in a country where ZIKV has been newly introduced. To study the dynamics of ZIKV early in an outbreak, we begin with human and vector populations at the disease-free equilibrium, and then introduce a single infected mosquito, following the time course of ZIKV transmission, including infections in pregnant women. To study dynamics of endemic ZIKV, we use a similar approach, but solve equations (1 – 6) numerically over a period of 500 years. This is sufficient time for the system to reach equilibrium across all parameter ranges considered, making it analogous to endemic spread.

### Sensitivity Analysis

We use a Latin Hypercube Sampling (LHS) scheme to explore model behavior over the full parameter space of our system. Specifically, we generate LHS matrices based on the parameter ranges in Table S1, and then use a Partial Rank Correlation Coefficient (PRCC) analysis to investigate the dependence of key system predictions (*R*_0_, endemic prenatal ZIKV exposures, peak prenatal ZIKV exposures, and endemic levels of immunity in the pre-reproductive class) on individual model parameters. For our LHS analysis of *R*_0_, we assume the full parameter ranges defined in Table S1. However, because the non-monotonicity of ZIKV exposures near *R*_0_ = 1 (see Figure 4 and Supplemental Information VI, Figure S7) would violate PRCC assumptions, for our LHS analysis of ZIKV exposures and immunity (where we use immunity to explain exposure patterns) we restrict the range for mosquito biting rates to *b* ∈ (0.4,1), for mosquito recruitment to *A* ∈ (1000, 5000) and for mosquito-to-human transmission to *β*_*h*_, ∈ (0.15, 0.75). This means that our PRCC analysis implicitly assumes that none of these parameters is so low that ZIKV spread is or is close to being unfavorable (i.e., *R*_0_ < 1). For all analyses, we use 10,000 samples and a uniform distribution across parameter ranges (notice this means that we use uniform distributions across *η*_*j*_, *η*_*xy*_, *η*_*m*_, *δ*, *μ* and *r* ranges, even though these parameters are reported as inverse values in Table S1 for ease of interpretation). We only consider 11 parameters, rather than the full 12 in Table S1, because we always fix the number of children per couple at two to ensure a constant human population size. This is done to simplify model analysis. In reality, most global populations, and particularly those in regions with ZIKV outbreaks, are growing. We explore the effect that a changing population size has on model predictions in Supplementary Information V.

